# A generic pipeline for CADD Score generation: chickenCADD and turkeyCADD

**DOI:** 10.1101/2024.11.01.621569

**Authors:** K. Lensing, JGC. van Schipstal, D. de Ridder, MAM. Groenen, MFL. Derks

## Abstract

Combined Annotation Dependent Depletion (CADD) is a machine learning approach used to predict the deleteriousness of genetic variants across a genome. By integrating diverse genomic features, CADD assigns a PHRED-like rank score to each potential variant. Unlike other methods, CADD does not rely on limited datasets of known pathogenic or benign variants but uses larger and less biased training sets. The rapid increase in high-quality genomes and functional annotations across species highlights the need for an automated, non-species-specific pipeline to generate CADD scores. Here, we introduce such a pipeline, facilitating the generation of CADD scores for various species using only a high-quality genome with gene annotation and a multi-species alignment. Additionally, we present updated chickenCADD scores and newly generated turkeyCADD scores, both generated with the pipeline.

## Introduction

Combined annotation dependent depletion (CADD) is an approach to predict the deleteriousness of genetic variants by machine learning. CADD was developed to rank variants, including single nucleotide variants (SNVs) and short insertions and deletions (InDels), throughout the human genome based on diverse genomic features derived from surrounding sequence context, gene model annotations, evolutionary constraint, epigenetic data and functional predictions (Rentzsch et al., 2019).The CADD methodology calculates a score for every possible single nucleotide variant (all three non-reference alleles), at every position of the genome via a machine learning model. All scores are then transformed into a PHRED-like rank score for improved interpretability (Groß, et al., 2020; Rentzsch et al., 2019).

Instead of using a relatively limited number of genomic variants for which pathogenic or benign status is ‘known’, CADD is trained on less biased and larger training sets. It relies on evolutional inference: variants that are fixed since a shared common ancestor with another (closely related) species, or those that are found in the population at a high frequency, are expected to be mostly benign. A second set of variants is simulated rather than derived from an inferred ancestor. These variants, free of selective pressure, are expected to be enriched for deleterious variants (Groß, et al., 2020; Kircher et al., 2014; Rentzsch et al., 2019). A machine learning model is trained to discriminate between these proxy-benign/neutral and proxy-deleterious variant classes.

CADD was first developed for humans (hCADD) (Kircher et al., 2014) and later adopted for mouse (mCADD) (Groß et al., 2018), chicken (chCADD) (Groß, et al., 2020) and pig (pCADD) (Groß, et al., 2020). The increasing availability of high-quality genomes and functional annotations call for the development of an automated generic pipeline to generate CADD scores for various species. Such a pipeline can also be used to easily update existing CADD scores based on new reference assemblies or newly available annotations, enabling more effective linkage of genotypic variation to specific traits.

Here we present a non-species-specific, automated and configurable pipeline to generate CADD scores. The pipeline integrates the CADD methodology to create scores with respect to their deleteriousness in the species genome. Additionally, we improved chicken, and generated turkey, CADD scores based on the latest reference genomes and annotations.

## Methods

The overall CADD-workflow consists of (1) the extraction of an inferred ancestral sequence, (2) generating variants, (3) annotating variants, (4) training a CADD model and (5) generating whole genome CADD scores (Figure 1). The pipeline was built using Snakemake (Mölder et al., 2021) based on the scripts provided by pCADD (Groß, et al., 2020) and chCADD (Groß, Bortoluzzi, et al., 2020). For a basic CADD model, the pipeline requires only a high-quality genome with gene annotation, and a multiple sequence alignment (MSA) containing the genome along with its phylogenetic tree which can be directly obtained from Ensembl Compara (Harrison et al., 2024). Additional genomic annotations and population frequency data can be added to the configuration file to build a more elaborate CADD model.

**Figure 1.**
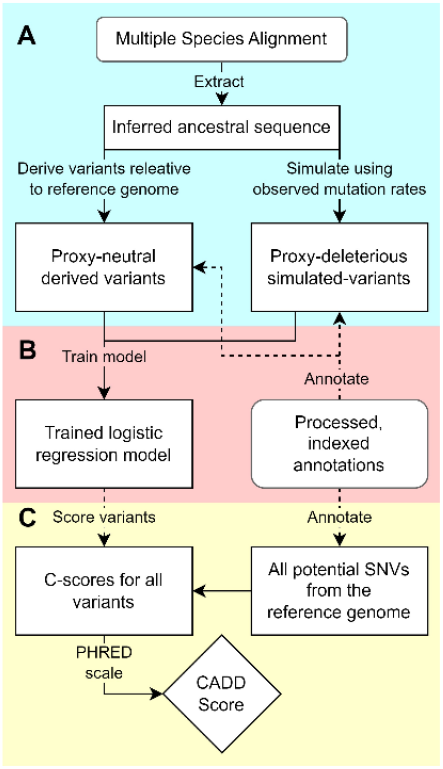
Overview of the devised CADD workflow. The approach consists of three key phases: A) Variant generation: proxy-neutral variants are derived from the inferred ancestral genome relative to the reference assembly, proxy-deleterious variants are simulated based on mutation rates found between the inferred ancestor and the reference assembly. B) A CADD model is trained on an equal number of proxy-neutral and proxy-deleterious variants represented by several provided and calculated annotations. C) Finally, all possible SNVs are annotated and scored by the trained model. By PHRED-scaling these scores the CADD scores are obtained.

### Multiple species alignment and inferred ancestral sequence

The first step of the pipeline extracts the inferred sequence of the most recent common ancestor of the species of interest and another configurable species present in the input MSA. Since the latest turkey reference assembly MGAL_WUR_HG_1.0 (Barros et al., 2022) is not (yet) included in the 27 sauropsids alignment of Compara release 112, a new MSA of 4 species (chicken (bGalGal1.mat.broiler.GRCg7b), turkey (MGAL_WU_HG_1.0), Japanese quail (Coturnix_japonica_2.0) and duck (CAU_duck1.0)) was created using the Progressive Cactus (v2.2.0) pipeline (Armstrong et al., 2020) (https://github.com/harvardinformatics/GenomeAnnotation-WholeGenomeAlignment) with default parameters and using hal2maf --chunkSize 1000000 -- filterGapCausingDupes --dupeMode single. Based on this MSA an inferred ancestral sequence was constructed for the species chicken and turkey.

### Derived and simulated variants

The set of derived variants containing proxy-benign/neutral variants comprises (nearly) fixed alleles that differ between the species genome and the inferred ancestral sequence. These fixed alleles have an allele frequency higher than 90% in the species population. Proxy-deleterious variants are simulated using local mutation rates inferred from the MSA. Where in mCADD, pCADD and chCADD these were derived from several ancestral sequences, the pipeline only uses rates between the reference genome and the ancestral sequence. For the new chCADD and tCADD, simulated variants observed in the population with allele frequencies higher than 10% were excluded, as they are likely not deleterious. Only variants found at sites that align with the ancestral genome were considered and an equal number of simulated variants were generated to match the derived variants. In total 23,444,020 proxy-benign/neutral SNVs were derived for chicken and 34,077,227 proxy-benign/neutral SNVs for turkey.

### Variant annotation

Different basic genomic annotations are automatically generated by the pipeline from the Ensembl Variant Effect Predictor database (McLaren et al., 2016) and can be supplemented with PhyloP scores (Pollard et al., 2010), PhastCons scores (Siepel et al., 2005), GERP conservation scores (Davydov et al., 2010), Grantham amino-acid substitution scores (Grantham, 1974), predictions of secondary DNA structure (Zhou et al., 2013) and functional annotations. The pipeline also automates the process of generating phastCons and phyloP conservation scores, which were found relevant in previous CADD studies (Groß, et al., 2020; Groß, et al., 2020; Groß et al., 2018). Annotations and their imputation for missing values can easily be added to the CADD model by the configuration file. Based on user configuration, categorical variables are one-hot encoded and missing values are handled by either a missing value indicator column, imputation of the population mean, or a fixed value.

A complete overview of the annotations used in the chCADD and tCADD model can be found in Tables S1 and S2. Missing values were imputed with the genome average obtained from the simulated data or set to 0 and all categorical values were recoded to binary variables. Basic annotations were obtained from VEP v110. Since SIFT scores (Ng & Henikoff, 2001) were not included in the VEP v110 annotations for turkey, they were computed separately using SIFT4G_Create_Genomic_DB and SIFT4G_Annotator (v. 2.0.0) (Vaser et al., 2015) based on the mgal_WUR_HG_1.0 assembly and annotation file. PhyloP and PhastCons scores were based on the 4-species MSA, excluding either chicken or turkey to avoid bias. GERP scores for chCADD were downloaded from Ensembl’s (release 112) extended version of the 27 sauropsids alignment. For tCADD, the GERP scores of Turkey_5.1 from the same 27 sauropsids alignment were lifted with Crossmap v0.6.6 (Zhao et al., 2014) to mgal_WUR_HG_1.0. Additionally, for chCADD chromatin states of 4 different tissues (muscle, lung, cortex and liver) from the FAANG project (Pan et al., 2023) were lifted from Ggal6 to GRCg7b with Crossmap and added to the model.

### Logistic regression model and scoring

The pipeline runs a logistic regression classifier (Scikitlearn (v1.3.1) (Pedregosa et al., 2011)) to distinguish between proxy-benign/neutral and proxy-deleterious variants based on the annotations added. The training parameters can be set using the configuration file. For chCADD and tCADD, the model was trained with a maximum of 100 iterations, the L2-penalization was set to 0.1 and the performance was assessed using 5-fold cross validation. Each possible SNV (3 per position) was annotated and scored by the CADD approach. Scores were sorted and assigned a CADD score defined as -10*log10(i/N), with i the rank of a SNV and N the total number of variants. ROC-AUC scores on the x-fold (x being configurable) cross-validated model are reported by the pipeline and additional validation on databases of experimentally known functional variants is included. However, as for chicken and turkey these databases were not available, performance is based on ROC-AUC scores.

## Results and discussion

### Pipeline

We developed a generic pipeline to obtain CADD scores for any species of interest. The Snakemake pipeline is a flexible and configurable workflow for training and validating CADD models and scoring variants. It is easy configurable with settings provided in a configuration file. This setup enables easy replication of the work, allows the workflow to be reused for new assemblies, and simplifies the addition of new or updated annotations to the CADD model.

### ChickenCADD and TurkeyCADD

The pipeline was used to generate chCADD and tCADD scores. The pipeline’s standard input is a MSA from Ensembl Compara. Because the new MGAL_WU_HG_1.0 assembly is not included in this MSA, a MSA consisting of four species (broiler, turkey, japanese quail and duck) was generated instead. This resulted in a high-quality ancestral sequence with high coverage between the species assembly and the inferred ancestral sequence (Figure S2).

A logistic regression model was trained to differentiate between two classes of variants, one set of putative deleterious simulated *de novo* variants and one set of putatively benign/neutral variants. The total number of SNVs used for model training was ∼47 million for chCADD and ∼68 million for tCADD. The top 10 model features with the largest weight in the fitted model are shown in Tables S4 and S5, and highlight the importance of conservation scores in CADD models. The performance of chCADD has improved considerably compared to the original ChCADD scores of Groß, et al. (2020). The mean ROC-AUC performance of the 5-fold cross-validated chCADD model is 0.805 (Figure 2), an improvement over the previous chCADD version’s ROC-AUC of 0.68. This improvement is likely due to a higher quality chicken genome, a high-quality ancestral sequence with high coverage between the chicken genome and the inferred ancestral sequence, and updated GERP scores. The high-coverage ancestral genome resulted in a larger and more accurate train-test set of variants, and the GERP scores are among the most important features in this and other animal CADD models (Groß et al., 2018, Groß et al., 2020). The mean ROC-AUC performance of the 5-fold cross-validated tCADD model is 0.728. The conversion of genomic coordinates for the GERP scores from meleagris_gallopavo.Turkey_5.1 to mgal_WUR_HG_1.0 likely explains the lower ROC-AUC, given the significant influence these scores had during model training. In general, both models seem to perform better on non-coding regions compared to the previous animal CADD models (Groß et al., 2018, Groß et al., 2020).

**Figure 2.**
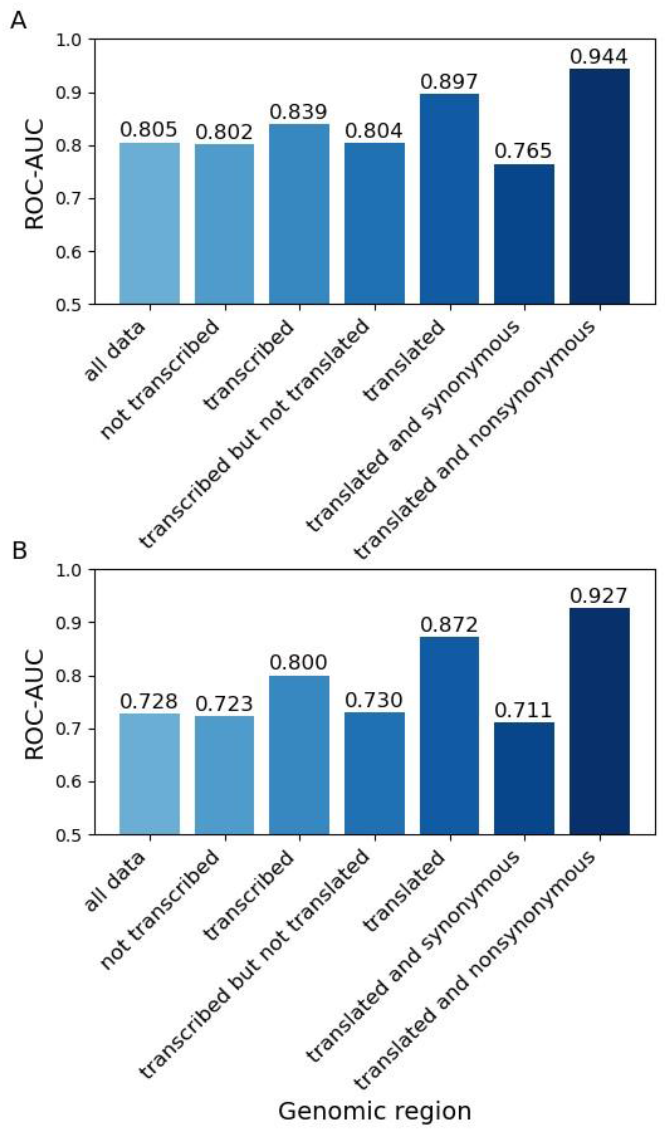
ROC-AUC scores on seven different subsets of the 5-fold cross-validated model reflecting different genomic regions and/or functional annotations of A. chCADD and B. tCADD.

The scores are scaled to PHRED-like scores as in the original CADD approach. Figure 3 shows the CADD scores for all possible variants in the chicken and turkey genome and their functional consequence as predicted by ENSEMBL VEP. Variants with a high CADD score are, as expected, annotated with “high impact” consequences such as stop-gained, canonical splice, and non-synonymous variants. Stop-gained variants were also the highest scoring in hCADD (Kircher et al., 2014).

**Figure 3.**
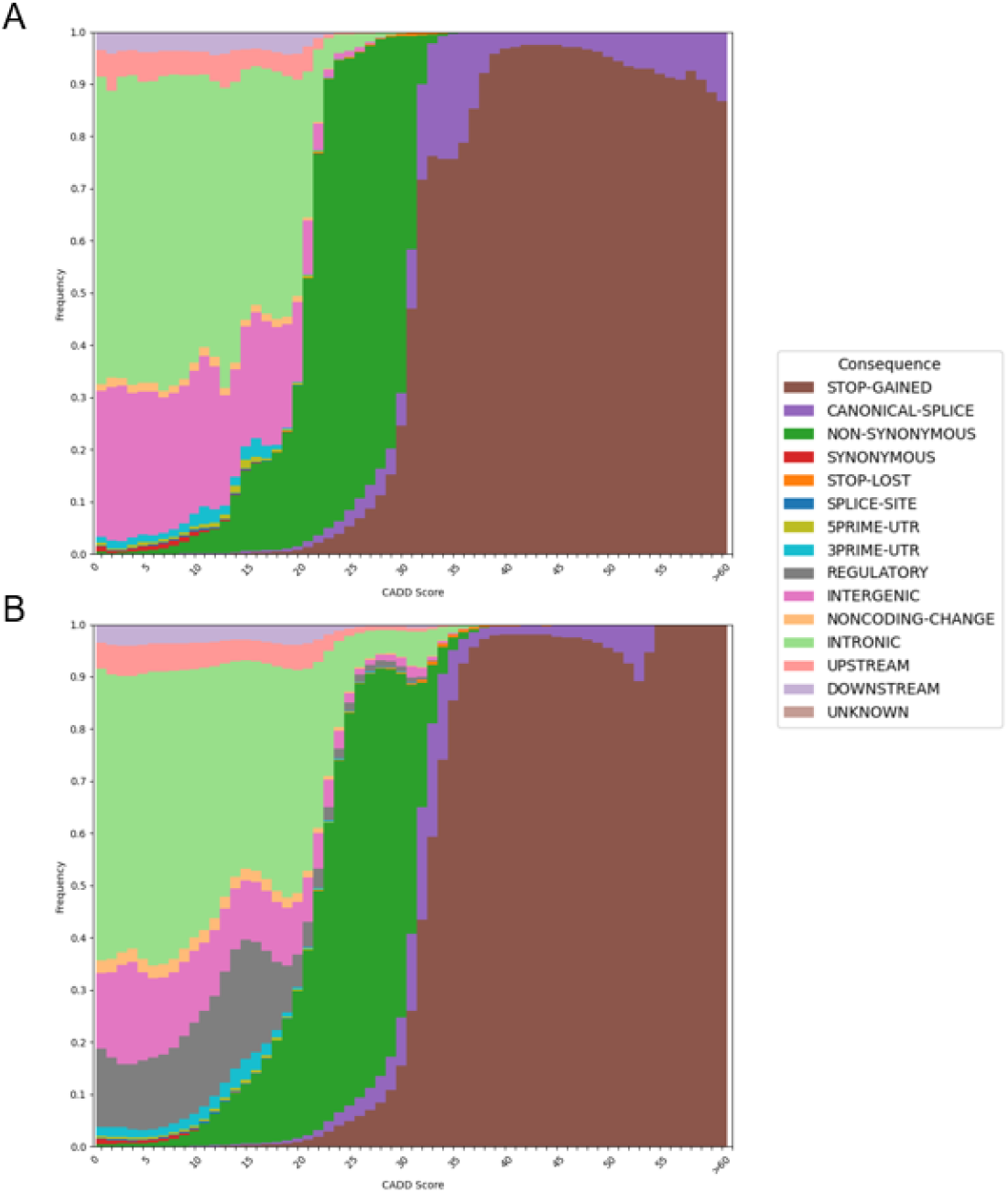
Scaled CADD scores of all A) chCADD and B) tCADD variants and frequency of assigned categorical variant consequences (Ensembl VEP).

## Conclusion

The automated CADD pipeline provides a robust, species-independent approach for generating CADD scores, as demonstrated by the improved chicken and newly developed turkey CADD scores. With substantial improvements in model performances, the pipeline represents a significant advancement in genomic variant scoring for livestock species. The generated scores are a useful resource to prioritize variants for further studies.

## Data Availability

The CADD pipeline is available from the WUR GitLab instance: https://git.wur.nl/job.vanschipstal/cadd-pipeline-v-2/-/tree/main?ref_type=heads

The chickenCADD and turkeyCADD scores can be downloaded from: https://public.anunna.wur.nl/ABGC/chickenCADD_v2/ and https://public.anunna.wur.nl/ABGC/turkeyCADD/

## Acknowledgment

We would like to thank Christian Groß for extensive feedback on his scripts and the CADD methodology and Seyan Hu for making a start with the pipeline.

## Supplemental data

**Figure S1.**
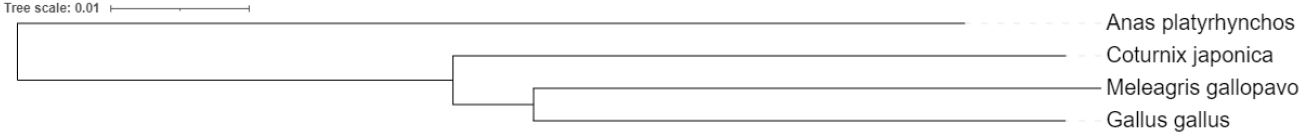
Phylogenetic tree used for creating the 4-species multiple sequence alignment with Progressive Cactus.

**Table S1.** List of annotations used for chCADD. Missing values are imputed via the specified values.

| Annotation label | Data type | Imputed value | Annotation description |
| --- | --- | --- | --- |
| Chrom | factor | - |  |
| Pos | int | - |  |
| Ref | factor | - | Reference allele |
| Alt | factor | - | Observed allele |
| isTv | bool | 0.5 | Is transversion? |
| Consequence | factor | - | VEP Consequence summaries |
| GC | num | 0.4 | Percent GC in a window of +/- 75bp |
| CpG | num | 0.02 | Percent CpG in a window of +/- 75bp |
| motifECount | int | 0 | Total number of overlapping motifs |
| motifEHIPos | factor | False | Is the position considered highly informative for an overlapping motif by VEP |
| motifEScoreChng | num | 0 | VEP score change for the overlapping motif site |
| Domain | factor | UD | Domain annotation inferred from VEP annotation (ncoils, tmhmm, sigp, lcompl, ndomain) |
| oAA | factor | UD | Amino acid of observed variant |
| nAA | factor | UD | Reference amino acid |
| Grantham | num | 0 | Grantham score: oAA,nAA |
| SIFTcat | factor | UD | SIFT category of change |
| SIFTval | num | 0 | SIFT score |
| cDNApos | num | 0 | Base position from transcription start |
| relcDNApos | num | 0 | Relative position in transcript |
| CDSpos | int | 0 | Base position from coding start |
| relCDSpos | num | 0 | Relative position in coding sequence |
| protPos | num | 0 | Amino acid position from coding start |
| relProtPos | num | 0 | Relative position in protein codon |
| dnaRoll | num | 0.2 | Predicted local DNA structure effect on dnaRoll |
| dnaProT | num | 0.5 | Predicted local DNA structure effect on dnaProT |
| dnaMGW | num | 0.01 | Predicted local DNA structure effect on dnaMGW |
| dnaHelT | num | -0.1 | Predicted local DNA structure effect on dnaHelT |
| GerpS | num | -0.3 | Rejected Substitution' score defined by GERP++ |
| verPhyloP | num | 0.04 | 4-msa PhastCons score (excl. chicken) |
| verPhCons | num | 0.2 | 4-msa PhyloP score (excl. chicken) |
| minDistTSS | num | 100000 | Distance to closest Transcribed Sequence Start (TSS) |
| minDistTSE | num | 100000 | Distance to closest Transcribed Sequence End (TSE) |
| Chrom-muscle | factor | UD | 15 distinct chromatin states liftover from galGal6 |
| Chrom-lung | factor | UD | 15 distinct chromatin states liftover from galGal6 |
| Chrom-cortex | factor | UD | 15 distinct chromatin states liftover from galGal6 |
| Chrom-liver | factor | UD | 15 distinct chromatin states liftover from galGal6 |

**Table S2.**
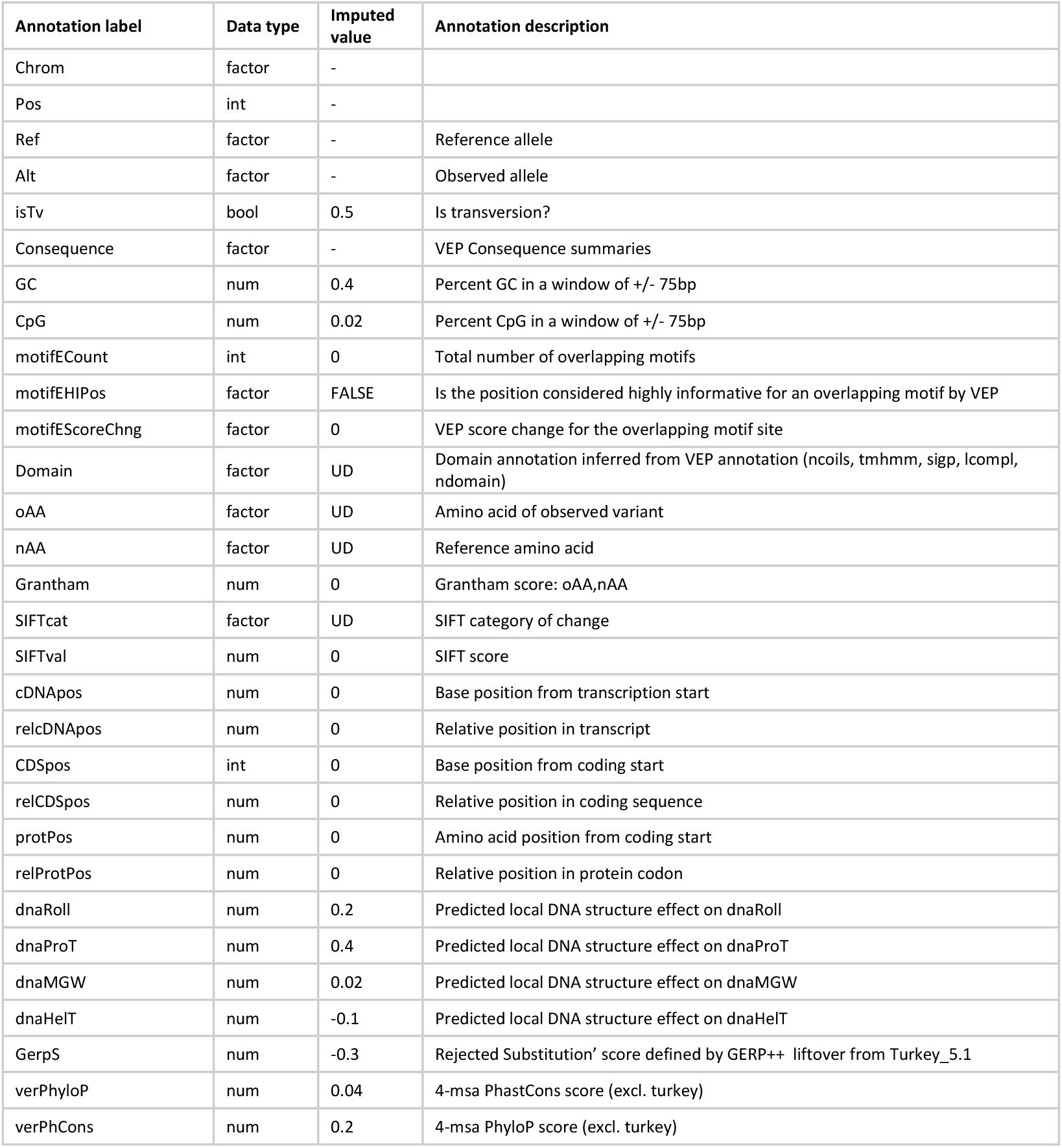
List of annotations used for tCADD. Missing values are imputed via the specified values.

**Table S3.** VEP consequences summarized in 14 categories. If a variant has multiple annotations, the consequence is selected based on the shown hierarchy.

| Hierarchy | Abbreviation | VEP Consequence |
| --- | --- | --- |
| 1 | SG | Stop Gained |
| 2 | CS | Canonical Splice |
| 3 | NS | Non-Synonymous |
| 4 | SN | Synonymous |
| 5 | SL | STOP Lost |
| 6 | S | Splice Site |
| 7 | U5 | 5'-UTR |
| 8 | U3 | 3'-UTR |
| 9 | IG | Intergenic |
| 10 | NC | Noncoding-change |
| 11 | I | Intronic |
| 12 | UP | Upstream |
| 13 | DN | Downstream |
| 14 | O | Unknown / Other |

**Figure S2.**
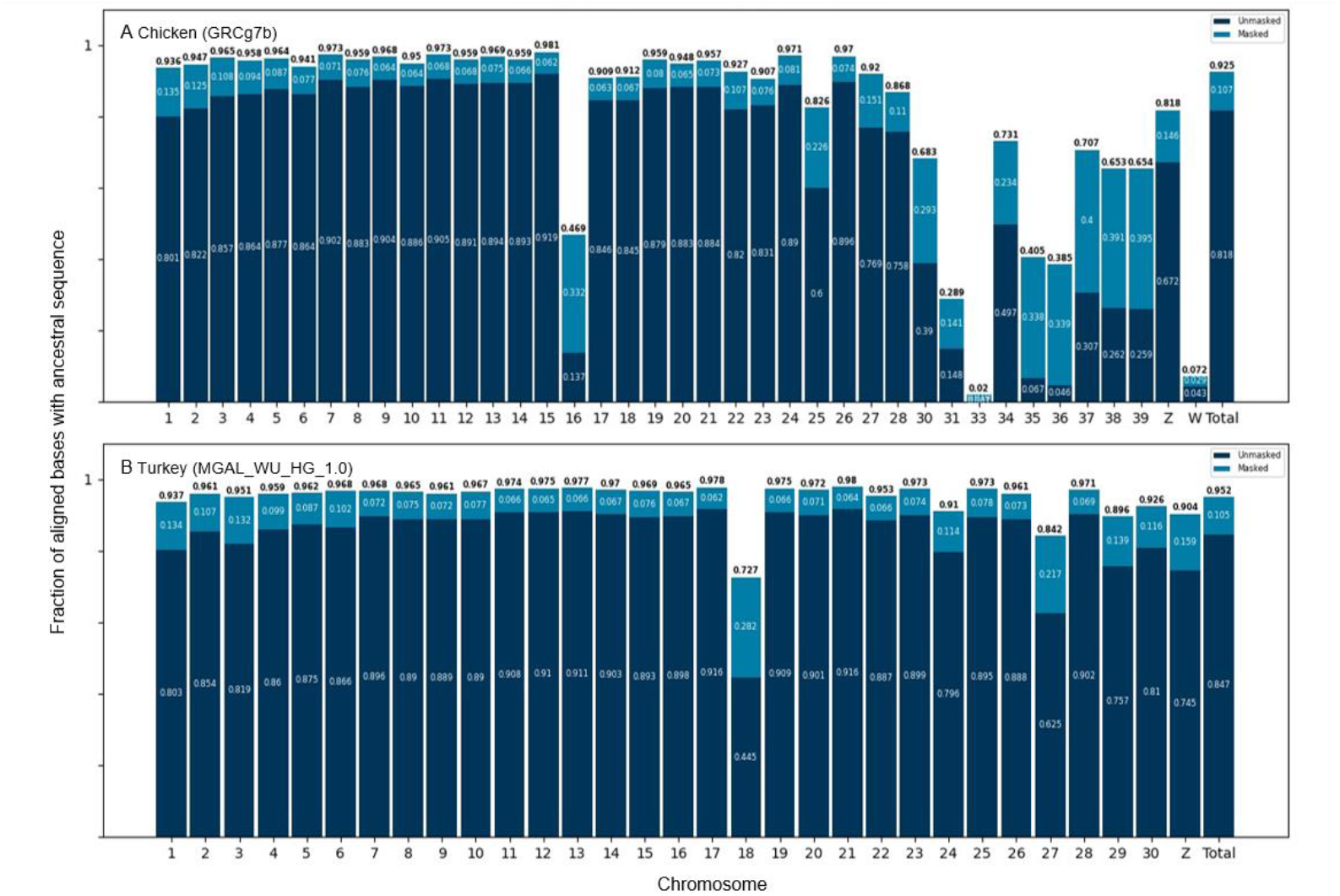
Coverage between A. the chicken bGalGal1.mat.broiler.GRCg7b assembly and the inferred ancestral sequence and B. the turkey MGAL_WU_HG_1.0 assembly and the inferred ancestral sequence. With low coverage for the micro chromosomes.

**Table S4.**
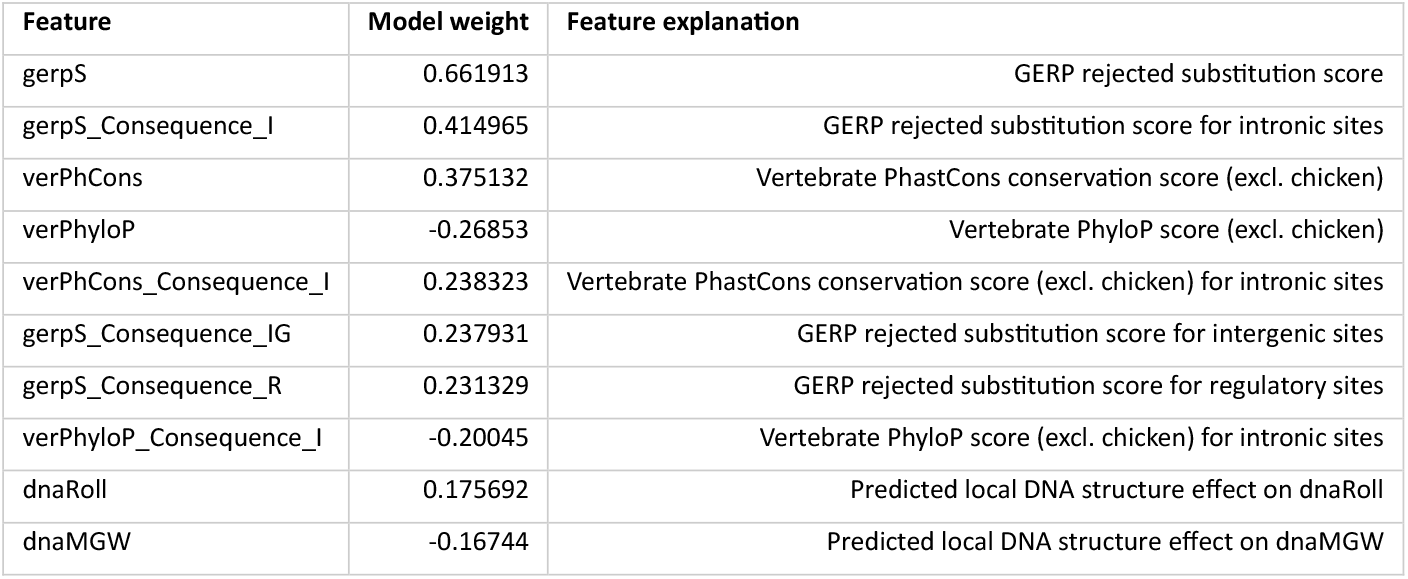
Top 10 model features in chickenCADD model with the largest assigned weight.

| Feature | Model weight | Feature explanation |
| --- | --- | --- |
| gerpS | 0.661913 | GERP rejected substitution score |
| gerpS_Consequence_I | 0.414965 | GERP rejected substitution score for intronic sites |
| verPhCons | 0.375132 | Vertebrate PhastCons conservation score (excl. chicken) |
| verPhyloP | -0.26853 | Vertebrate PhyloP score (excl. chicken) |
| verPhCons_Consequence_I | 0.238323 | Vertebrate PhastCons conservation score (excl. chicken) for intronic sites |
| gerpS_Consequence_IG | 0.237931 | GERP rejected substitution score for intergenic sites |
| gerpS_Consequence_R | 0.231329 | GERP rejected substitution score for regulatory sites |
| verPhyloP_Consequence_I | -0.20045 | Vertebrate PhyloP score (excl. chicken) for intronic sites |
| dnaRoll | 0.175692 | Predicted local DNA structure effect on dnaRoll |
| dnaMGW | -0.16744 | Predicted local DNA structure effect on dnaMGW |

**Table S5.** Top 10 model features in turkeyCADD model with the largest assigned weight.

| Feature | Model weight | Feature explanation |
| --- | --- | --- |
| GerpS | 0.381656 | GERP rejected substitution score |
| verPhCons | 0.353878 | Vertebrate PhastCons conservation score (excl. turkey) |
| GerpS_Consequence_I | 0.254803 | GERP rejected substitution score for intronic sites |
| verPhCons_Consequence_I | 0.228232 | Vertebrate PhastCons conservation score (excl. turkey) for intronic sites |
| GerpS_Consequence_IG | 0.18503 | GERP rejected substitution score for intergenic sites |
| SIFTval | -0.18453 | SIFT score |
| verPhCons_Consequence_IG | 0.148624 | Vertebrate PhastCons conservation score (excl. turkey) for intergenic sites |
| SIFTcat_deleterious | 0.134344 | SIFT category of change deleterious |
| verPhCons_Consequence_NS | 0.118971 | Vertebrate PhastCons conservation score (excl. turkey) for non-synonymous sites |
| GerpS_Consequence_UP | 0.090974 | GERP rejected substitution score for upstream sites |

**Table S6.**
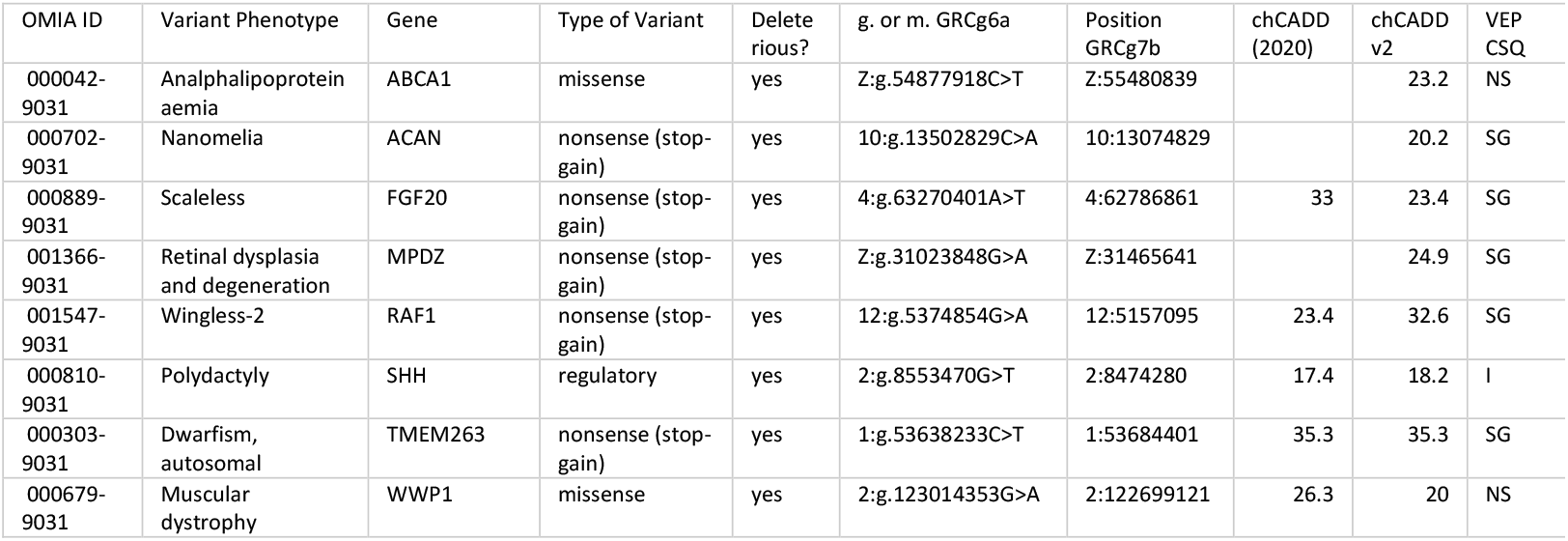
Likely causal variants of phenotype changes from the Online Mendelian Inheritance in Animals (OMIA) database and their CADD scores. Location of variants were mapped from GRCg6a reference genome to GRCg7b via CrossMap.

| OMIA ID | Variant Phenotype | Gene | Type of Variant | Deleterious? | g. or m. GRCg6a | Position GRCg7b | chCADD (2020) | chCADD v2 | VEP CSQ |
| --- | --- | --- | --- | --- | --- | --- | --- | --- | --- |
| 000042-9031 | Analphalipoprotein aemia | ABCA1 | missense | yes | Z:g.54877918C>T | Z:55480839 |  | 23.2 | NS |
| 000702-9031 | Nanomelia | ACAN | nonsense (stop-gain) | yes | 10:g.13502829C>A | 10:13074829 |  | 20.2 | SG |
| 000889-9031 | Scaleless | FGF20 | nonsense (stop-gain) | yes | 4:g.63270401A>T | 4:62786861 | 33 | 23.4 | SG |
| 001366-9031 | Retinal dysplasia and degeneration | MPDZ | nonsense (stop-gain) | yes | Z:g.31023848G>A | Z:31465641 |  | 24.9 | SG |
| 001547-9031 | Wingless-2 | RAF1 | nonsense (stop-gain) | yes | 12:g.5374854G>A | 12:5157095 | 23.4 | 32.6 | SG |
| 000810-9031 | Polydactyly | SHH | regulatory | yes | 2:g.8553470G>T | 2:8474280 | 17.4 | 18.2 | I |
| 000303-9031 | Dwarfism, autosomal | TMEM263 | nonsense (stop-gain) | yes | 1:g.53638233C>T | 1:53684401 | 35.3 | 35.3 | SG |
| 000679-9031 | Muscular dystrophy | WWP1 | missense | yes | 2:g.123014353G>A | 2:122699121 | 26.3 | 20 | NS |

## Literature cited

Armstrong, J., Hickey, G., Diekhans, M., Fiddes, I. T., Novak, A. M., Deran, A., Fang, Q., Xie, D., Feng, S., Stiller, J., Genereux, D., Johnson, J., Marinescu, V. D., Alföldi, J., Harris, R. S., Lindblad-Toh, K., Haussler, D., Karlsson, E., Jarvis, E. D., … Paten, B. (2020). Progressive Cactus is a multiple-genome aligner for the thousand-genome era. Nature 2020 587:7833, 587(7833), 246–251. 10.1038/s41586-020-2871-y

Barros, C. P., Derks, M. F. L., Mohr, J., Wood, B. J., Crooijmans, R. P. M. A., Megens, H. J., Bink, M. C. A. M., & Groenen, M. A. M. (2022). A new haplotype-resolved turkey genome to enable turkey genetics and genomics research. GigaScience, 12, 1–14. 10.1093/GIGASCIENCE/GIAD051

Davydov, E. V., Goode, D. L., Sirota, M., Cooper, G. M., Sidow, A., & Batzoglou, S. (2010). Identifying a High Fraction of the Human Genome to be under Selective Constraint Using GERP++. PLOS Computational Biology, 6(12), e1001025. 10.1371/JOURNAL.PCBI.1001025

GitHub - harvardinformatics/GenomeAnnotation-WholeGenomeAlignment: guidelines for running progressive CACTUS to produce a whole genome alignment required for genome annotation tools that leverage WGA. (n.d.). Retrieved July 23, 2024, from https://github.com/harvardinformatics/GenomeAnnotation-WholeGenomeAlignment

Grantham, R. (1974). Amino Acid Difference Formula to Help Explain Protein Evolution. Science, 185(4154), 862–864. 10.1126/SCIENCE.185.4154.862

Groß, C., Bortoluzzi, C., Ridder, D. de, Megens, H. J., Groenen, M. A. M., Reinders, M., & Bosse, M. (2020). Prioritizing sequence variants in conserved non-coding elements in the chicken genome using chCADD. PLOS Genetics, 16(9), e1009027. 10.1371/JOURNAL.PGEN.1009027

Groß, C., de Ridder, D., & Reinders, M. (2018). Predicting variant deleteriousness in non-human species: Applying the CADD approach in mouse. BMC Bioinformatics, 19(1), 1–10. 10.1186/S12859-018-2337-5/FIGURES/5

Groß, C., Derks, M., Megens, H. J., Bosse, M., Groenen, M. A. M., Reinders, M., & De Ridder, D. (2020). PCADD: SNV prioritisation in Sus scrofa. Genetics Selection Evolution, 52(1), 1–15. 10.1186/S12711-020-0528-9/TABLES/5

Harrison, P. W., Amode, M. R., Austine-Orimoloye, O., Azov, A. G., Barba, M., Barnes, I., Becker, A., Bennett, R., Berry, A., Bhai, J., Bhurji, S. K., Boddu, S., Lins, P. R. B., Brooks, L., Ramaraju, S. B., Campbell, L. I., Martinez, M. C., Charkhchi, M., Chougule, K., … Yates, A. D. (2024). Ensembl 2024. Nucleic Acids Research, 52(D1), D891–D899. 10.1093/NAR/GKAD1049

Kircher, M., Witten, D. M., Jain, P., O’roak, B. J., Cooper, G. M., & Shendure, J. (2014). A general framework for estimating the relative pathogenicity of human genetic variants. Nature Genetics 2014 46:3, 46(3), 310–315. 10.1038/ng.2892

McLaren, W., Gil, L., Hunt, S. E., Riat, H. S., Ritchie, G. R. S., Thormann, A., Flicek, P., & Cunningham, F. (2016). The Ensembl Variant Effect Predictor. Genome Biology, 17(1), 1–14. 10.1186/S13059-016-0974-4/TABLES/8

Mölder, F., Jablonski, K. P., Letcher, B., Hall, M. B., Tomkins-Tinch, C. H., Sochat, V., Forster, J., Lee, S., Twardziok, S. O., Kanitz, A., Wilm, A., Holtgrewe, M., Rahmann, S., Nahnsen, S., & Köster, J. (2021). Sustainable data analysis with Snakemake. F1000Research 2021 10:33, 10, 33. 10.12688/f1000research.29032.1

Ng, P. C., & Henikoff, S. (2001). Predicting Deleterious Amino Acid Substitutions. Genome Research, 11(5), 863–874. 10.1101/GR.176601

Pan, Z., Wang, Y., Wang, M., Wang, Y., Zhu, X., Gu, S., Zhong, C., An, L., Shan, M., Damas, J., Halstead, M. M., Guan, D., Trakooljul, N., Wimmers, K., Bi, Y., Wu, S., Delany, M. E., Bai, X., Cheng, H. H., … Zhou, H. (2023). An atlas of regulatory elements in chicken: A resource for chicken genetics and genomics. Science Advances, 9(18). 10.1126/SCIADV.ADE1204/SUPPL_FILE/SCIADV.ADE1204_TABLES_S1_TO_S12.ZIP

Pedregosa FABIANPEDREGOSA, F., Michel, V., Grisel OLIVIERGRISEL, O., Blondel, M., Prettenhofer, P., Weiss, R., Vanderplas, J., Cournapeau, D., Pedregosa, F., Varoquaux, G., Gramfort, A., Thirion, B., Grisel, O., Dubourg, V., Passos, A., Brucher, M., Perrot andÉdouardand, M. Duchesnay, andÉdouard, & Duchesnay EDOUARDDUCHESNAY, Fré. (2011). Scikit-learn: Machine Learning in Python. Journal of Machine Learning Research, 12(85), 2825–2830. http://jmlr.org/papers/v12/pedregosa11a.html

Pollard, K. S., Hubisz, M. J., Rosenbloom, K. R., & Siepel, A. (2010). Detection of nonneutral substitution rates on mammalian phylogenies. Genome Research, 20(1), 110–121. 10.1101/GR.097857.109

Rentzsch, P., Witten, D., Cooper, G. M., Shendure, J., & Kircher, M. (2019). CADD: predicting the deleteriousness of variants throughout the human genome. Nucleic Acids Research, 47(D1), D886–D894. 10.1093/NAR/GKY1016

Siepel, A., Bejerano, G., Pedersen, J. S., Hinrichs, A. S., Hou, M., Rosenbloom, K., Clawson, H., Spieth, J., Hillier, L. D. W., Richards, S., Weinstock, G. M., Wilson, R. K., Gibbs, R. A., Kent, W. J., Miller, W., & Haussler, D. (2005). Evolutionarily conserved elements in vertebrate, insect, worm, and yeast genomes. Genome Research, 15(8), 1034–1050. 10.1101/GR.3715005

Vaser, R., Adusumalli, S., Ngak Leng, S., Sikic, M., & Ng, P. C. (2015). SIFT missense predictions for genomes. Nature Protocols, 11. 10.1038/nprot.2015.123

Zhao, H., Sun, Z., Wang, J., Huang, H., Kocher, J. P., & Wang, L. (2014). CrossMap: a versatile tool for coordinate conversion between genome assemblies. Bioinformatics (Oxford, England), 30(7), 1006–1007. 10.1093/BIOINFORMATICS/BTT730

Zhou, T., Yang, L., Lu, Y., Dror, I., Dantas Machado, A. C., Ghane, T., Di Felice, R., & Rohs, R. (2013). DNAshape: a method for the high-throughput prediction of DNA structural features on a genomic scale. Nucleic Acids Research, 41(W1), W56–W62. 10.1093/NAR/GKT437

